# Time-resolved network control analysis links reduced control energy under DMT with the serotonin 2a receptor, signal diversity, and subjective experience

**DOI:** 10.1101/2023.05.11.540409

**Authors:** S. Parker Singleton, Christopher Timmermann, Andrea I. Luppi, Emma Eckernäs, Leor Roseman, Robin L. Carhart-Harris, Amy Kuceyeski

**Author notes:** Corresponding author Email address (S. Parker Singleton).

## Abstract

Psychedelics offer a profound window into the functioning of the human brain and mind through their robust acute effects on perception, subjective experience, and brain activity patterns. In recent work using a receptor-informed network control theory framework, we demonstrated that the serotonergic psychedelics lysergic acid diethylamide (LSD) and psilocybin flatten the brain’s control energy landscape in a manner that covaries with more dynamic and entropic brain activity. Contrary to LSD and psilocybin, whose effects last for hours, the serotonergic psychedelic N,N-dimethyltryptamine (DMT) rapidly induces a profoundly immersive altered state of consciousness lasting less than 20 minutes, allowing for the entirety of the drug experience to be captured during a single resting-state fMRI scan. Using network control theory, which quantifies the amount of input necessary to drive transitions between functional brain states, we integrate brain structure and function to map the energy trajectories of 14 individuals undergoing fMRI during DMT and placebo. Consistent with previous work, we find that global control energy is reduced following injection with DMT compared to placebo. We additionally show longitudinal trajectories of global control energy correlate with longitudinal trajectories of EEG signal diversity (a measure of entropy) and subjective ratings of drug intensity. We interrogate these same relationships on a regional level and find that the spatial patterns of DMT’s effects on these metrics are correlated with serotonin 2a receptor density (obtained from separately acquired PET data). Using receptor distribution and pharmacokinetic information, we were able to successfully recapitulate the effects of DMT on global control energy trajectories, demonstrating a proof-of-concept for the use of control models in predicting pharmacological intervention effects on brain dynamics.

## Introduction

Serotonergic psychedelics such as lysergic acid diethylamide (LSD), psilocybin, and N,N-dimethyltrypamine (DMT) are powerful neuromodulators that transiently alter human experience (Nichols 2004; Shulgin and Shulgin 1997) and have shown potential for treating a variety of common affective and addictive disorders (Andersen et al. 2021; Vollenweider and Smallridge 2022). DMT is a naturally occurring tryptamine and is the primary psychoactive compound found in ayahuasca, a ceremonial brew used for hundreds of years in South America (Metzner 2005). Unlike LSD and psilocybin, DMT is rapidly metabolized in the body by mono-amine oxidase (MAO) enzymes, requiring it to be combined with MAO-inhibitors in order to be orally active, as is the case in ayahuasca. This results in a DMT experience that rises and falls over the course of several hours, similar to oral LSD and psilocybin. When inhaled or injected intravenously at large enough doses, however, DMT rapidly produces immersive “breakthrough” experiences characterized by vivid and complex visual imagery - occurring within one minute of administration - and lasting for only 15-30 minutes (Lawrence et al. 2022; Timmermann et al. 2019; Strassman et al. 1994; Strassman 1995). This provides a unique opportunity to study human brain dynamics during the onset, peak, and offset of DMT’s effects over a single functional scan.

Human neuroimaging studies with LSD and psilocybin have demonstrated that the psychedelic state is one of prominent reorganization of brain dynamics (Carhart-Harris and Friston 2019; Vollenweider and Preller 2020; Doss, Madden, et al. 2021; McCulloch et al. 2022). These compounds acutely decrease integrity within the brian’s functional sub-networks, while increasing integrity between functional sub-networks (Roseman et al. 2014; McCulloch et al. 2022; Girn et al. 2022; 2023; Dai et al. 2023). The impact of psychedelics on subjective experience (Kraehenmann et al. 2017), neural dynamics (Preller, Burt, et al. 2018; Preller, Schilbach, et al. 2018), and therapeutic behavior change (Cameron et al. 2023) has been linked to agonism of the serotonin 2a (5-HT2a) receptor. This finding affords a unique opportunity to model and study the perturbation of brain dynamics using whole-brain computational models that incorporate receptor distribution information. Such whole-brain models have highlighted the central role of the spatial distribution of the 5-HT2a receptor in the shift in brain dynamics under LSD and psilocybin (Deco et al. 2018; Kringelbach et al. 2020; Singleton et al. 2022).

Network control theory is a linear dynamical systems approach that models state transitions occurring within a network (Gu et al. 2015). When applied to the brain, typically the structural connectivity matrix derived from diffusion MRI (dMRI) is the network over which transitions between functional brain states are modeled. These functional brain states may be theoretical, e.g. activations of *a priori* functional networks (He et al. 2022; Parkes et al. 2022), or empirical, e.g. statistical brain maps derived from task functional MRI (fMRI) (Braun et al. 2021; Luppi et al. 2023) or commonly recurring patterns of co-activation derived from the clustering of task-free fMRI time-series (Cornblath et al. 2020; Singleton et al. 2022). Overall, network control theory has demonstrated utility in describing brain dynamics in a variety of cognitive states (Cornblath et al. 2020; Zhou et al. 2021) and neuropsychiatric/degenerative conditions (He et al. 2022; Braun et al. 2021; Parkes et al. 2021; Tozlu et al. 2023), as well as throughout development (Parkes et al. 2022; Cornblath et al. 2019) and during neuromodulation and pharmacologically induced altered states (Singleton et al. 2022; Stiso et al. 2019; Luppi et al. 2023).

Receptor-informed network control theory is an extension we recently deployed in order to model the effects of LSD and psilocybin on brain activity dynamics. We found that the acute administration of LSD and psilocybin reduces the control energy required to transition between task-free fMRI-derived brain-states in a manner that, across individuals, covaries with increases in brain activity entropy - i.e., the diversity or complexity of the brain’s spontaneous oscillations recorded across time, a well-known marker of psychedelic action (Carhart-Harris 2018). Moreover, we provided evidence that the enhanced state-transitioning effect of psychedelics is associated with the brain’s spatial distribution of the 5-HT2a receptor expression (Singleton et al. 2022). In that work, we studied the transitions between and dynamics of four representative activity patterns. While this approach is ideal for summarizing overall changes, it lacks the temporal resolution necessary for capturing instantaneous, moment-to-moment, shifts in dynamics under a rapidly changing cognitive state such as when under the influence of DMT.

Given the rapid kinetics of DMT’s effects, the use of time-resolved analysis techniques will be crucial for capturing changes in the brain’s activation dynamics in real time. Here, we employ a time-resolved network control analysis of N=14 healthy individuals undergoing simultaneous electroencephalography (EEG) and fMRI recordings for 8 minutes before and 20 minutes after an intravenous (I.V.) bolus injection of DMT and (on a separate visit) placebo (Figure 1a) (Timmermann et al. 2023). These multimodal and continuous scanning conditions enable high temporal (EEG) and spatial (fMRI) resolution of brain activity before, during, and after an injection of DMT. Herein, we expand upon our prior work with LSD and psilocybin (Singleton et al. 2022) by deploying a time-resolved network control analysis of the entire trajectory of the effects of DMT (Figure 1b). We compare control energy dynamics between DMT and placebo, observe temporal trajectories of these dynamics, and relate these changes to contemporaneous changes in neuronal signal diversity (Lempel-Ziv complexity; a measure approximating entropy) from concurrently-acquired EEG. Further, we compare DMT-related changes in regional dynamics to various serotonin receptor maps, including 2a. Lastly, we demonstrate an ability to simulate the impacts of DMT on control energy dynamics *in silico* using only fMRI data from placebo scans, 2a receptor density information, and pharmacokinetic modeling of DMT plasma concentrations.

**Figure 1:**
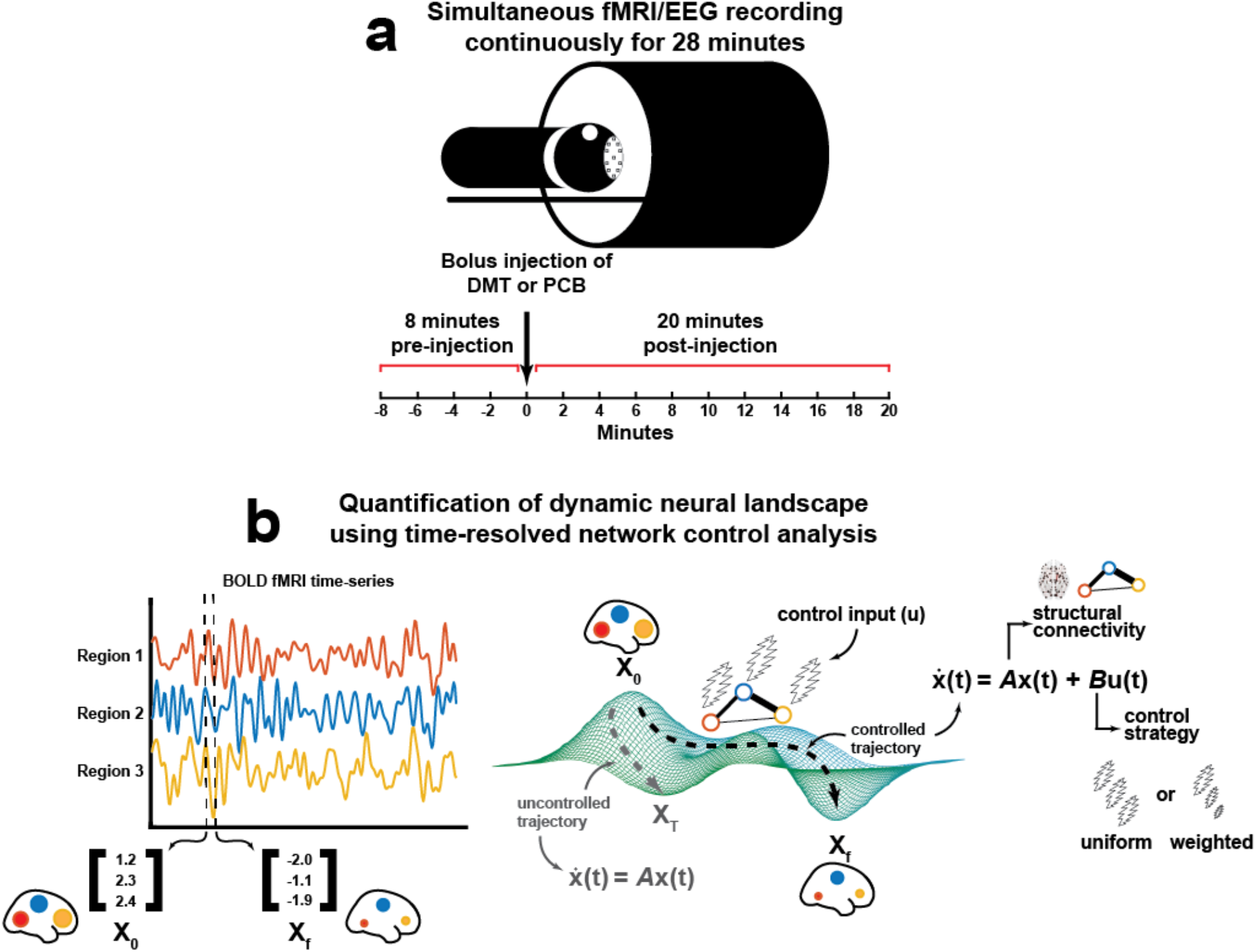
Time-resolved network control analysis of the human brain during a pharmacologically-induced alteration of consciousness. **(a)** Fourteen individuals were scanned over two days in which they received either DMT and saline placebo in separate visits (two-weeks apart, single-blind, counterbalanced design). On each day, a 28-minute long eyes-closed resting-state EEG-fMRI scan was performed with DMT/placebo intravenously administered at the end of the 8th minute. On the same day, identical scanning sessions were performed where participants were asked to rate the subjective intensity of drug effects at the end of every minute. **(b)** Here, we deploy a time-resolved network control analysis of the brain’s trajectory through its activational landscape. The position in the landscape is illustrated here as a 3D vector containing regional BOLD signal amplitude at a given time t. We compute a control energy time-series from the regional activity vector time series by modeling transitions between adjacent regional activity vectors (x_0_ and x_f_, respectively) using a linear time-invariant model within a network control theory framework. In this framework, the state of the network x(t), here a vector of regional BOLD activations at time t, evolves over time via diffusion through the brain’s weighted structural connectome *A*, the adjacency matrix. In order to complete the desired transition from the initial (x_0_) to the target state (x_f_), input (u) is injected into each region in the network. Varying control strategies (reflected in the matrix *B*) may be deployed wherein different regions are assigned varied amounts of control within the system. Integrating input u(t) at each node over the length of the trajectory from x_0_ to x_f_ yields region-wise control energy, and summing over all regions yields a global value of control energy required to complete the transition.

## Results

We analyzed simultaneous EEG-fMRI resting-state data for 14 participants acquired across two sessions, each on separate days (Timmermann et al. 2023). At each session, 28 minute long resting-state EEG-fMRI scans were collected, with I.V. bolus infusion of either DMT or placebo at the end of the 8th minute (Figure 1a).

### Global control energy is lower after DMT infusion versus after placebo

We first begin by computing a control energy time-series from each participant’s 28 minute resting-state DMT and placebo fMRI scans. Control energy here is defined as the amount of input needed to drive the system from the current brain activity pattern to the next, where each brain activity pattern is a regional vector summarizing a single brain volume within the fMRI (Figure 1b). We find that DMT control energies are significantly lower than placebo control energies for a majority of the time points (65.5%; corrected p < 0.05) in the 20 minutes following injection (Figure 2a shows group average time series).

**Figure 2:**
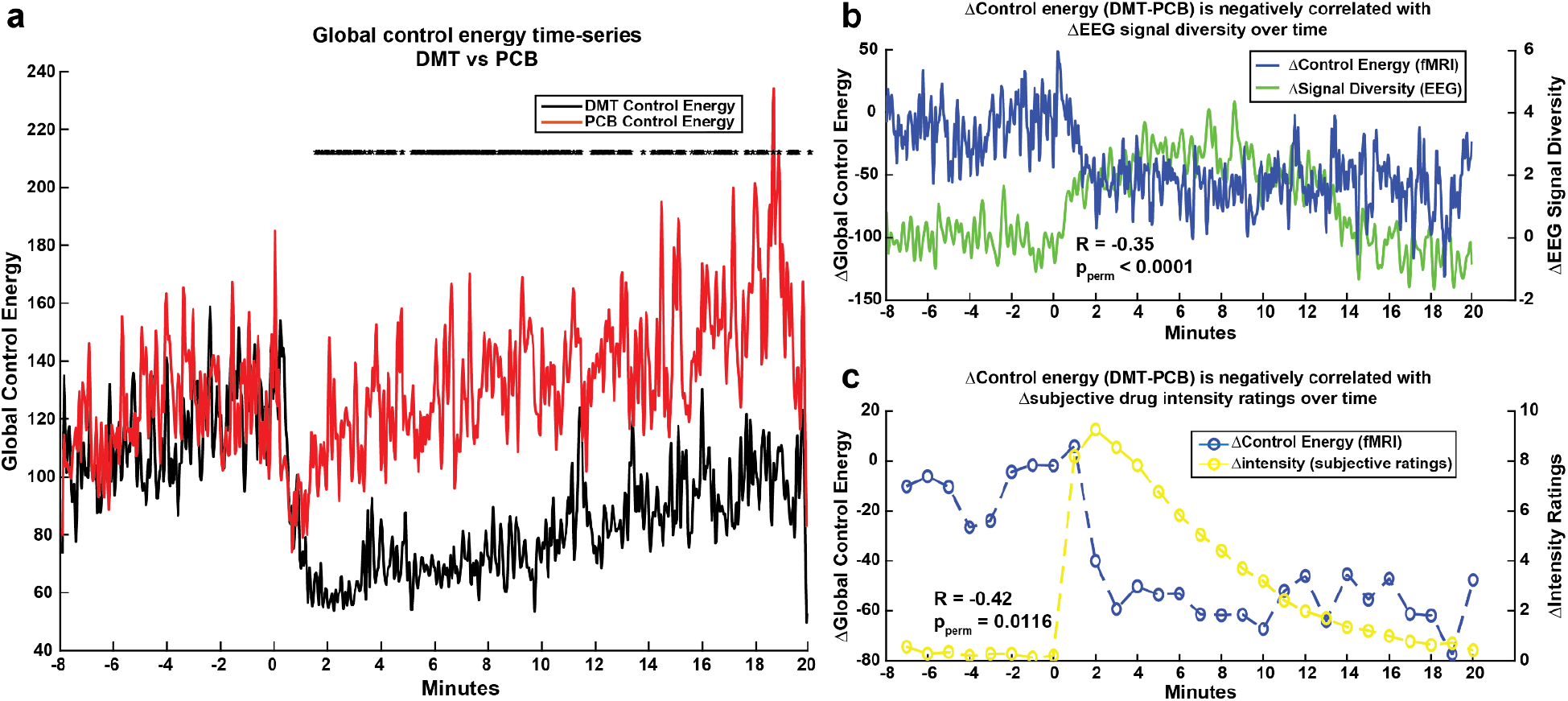
Global control energy is reduced after DMT injection compared to after placebo injection, and negatively correlates with signal diversity and subjective drug intensity ratings. **(a)** Group-average global control energy time-series for the DMT and placebo (PCB) conditions. Global control energies for each time point’s transition following injection were compared via a two-sided, paired t-test and p-values were corrected for multiple comparisons using the Benjamini-Hochberg method (Benjamini and Hochberg 1995). Nearly two-thirds of post-injection control energies (65.5%) were found to be significantly lower under DMT compared to placebo (* = corrected p < 0.05). **(b)** Differences in global control energy and EEG signal diversity between the DMT and PCB conditions are negatively correlated over the 28 minute scans (Pearson’s R = -0.348, p_perm_ < 0.0001), indicating that lower demand for fMRI-based global control energy was associated with increased EEG-based signal diversity of brain activity. **(c)** Differences in global control energy between the DMT and PCB conditions were averaged over one minute intervals in order to compare with subjective drug intensity ratings (0 -10) collected at the end of each minute (the latter of which were obtained during a separate fMRI from the one used to calculate global control energy). We found a negative correlation over time between intensity ratings and differences between the DMT and placebo conditions’ global control energies (Pearson’s R = -0.418, p_perm_ 0.0116).

### Global control energy under DMT negatively correlates with subjective drug intensity ratings and signal diversity from simultaneous EEG

We next relate the dynamical changes observed under DMT to signal diversity, quantified in terms of Lempel-Ziv complexity, derived from simultaneous EEG recordings. We correlate the between-condition differences in these metrics and find that the more fMRI-based global control energy is decreased under DMT, the more signal diversity from EEG is increased (Figure 2b; Pearson’s R = -0.34801, p_perm_ < 0.0001). In separate scanning sessions with an identical dosing regimen, the same participants rated the subjective intensity of drug effects on a scale of 0-10 at the end of every minute. We reduced the dimensionality of the control energy time-series by averaging over one-minute windows corresponding to the collection of intensity ratings. Correlating the between-condition differences in these metrics, we find that reduction of control energy by DMT correlates with the intensity of the drug’s subjective effects over time (Figure 2c; Pearson’s R = -0.4184, p_perm_ = 0.0116).

### Spatial patterns of control energy and its correlation with signal diversity and intensity are associated with spatial patterns of serotonin 2a receptors

Next, we interrogate control energy under DMT at the regional, rather than global, level. Regional control energy reflects the amount of input injected into each region in order to complete a desired transition, whereas global control energy is the sum of this metric over all regions. While global control energy provides a single metric useful for summarizing the relative difficulty of state transitions, regional control energy can help quantify the varied contributions of each brain region to the global measure. We evaluated regional control energy under DMT in three ways (Figure 3a): 1) the change in regional control energy post-injection relative to pre-injection, 2) each region’s control energy correlated with global EEG signal diversity over time during DMT scans, and 3) each region’s control energy correlated with subjective drug intensity over time during DMT scans. Reflecting the global result in Figure 2, we find that regional control energy is generally decreased following DMT injection, and is generally inversely correlated with both EEG signal diversity and drug intensity ratings (Figure 3a). Our purpose in assessing control energy at a regional level is to relate its spatial distribution to biologically relevant patterns in the human brain. To do so, we calculated the Spearman rank correlation between each of these regional metrics and the serotonin 2a receptor cortical density distribution derived from PET imaging (Figure 3b) (Beliveau et al. 2017). To assess the robustness of these regional metrics’ correlations with the 2a cortical map, we compared their values against 10,000 correlations calculated with spun permuted (Váša et al. 2018) 2a cortical maps that preserve spatial autocorrelations present in the original maps. We find that each of our regional metrics from Figure 3a is negatively correlated with the serotonin 2a spatial distribution. These negative correlations indicate that brain regions with more serotonin 2a receptors have their control energies decreased more by DMT, and their control energies over time are more negatively correlated with EEG signal diversity and subjective drug intensity (Figure 3c).

**Figure 3:**
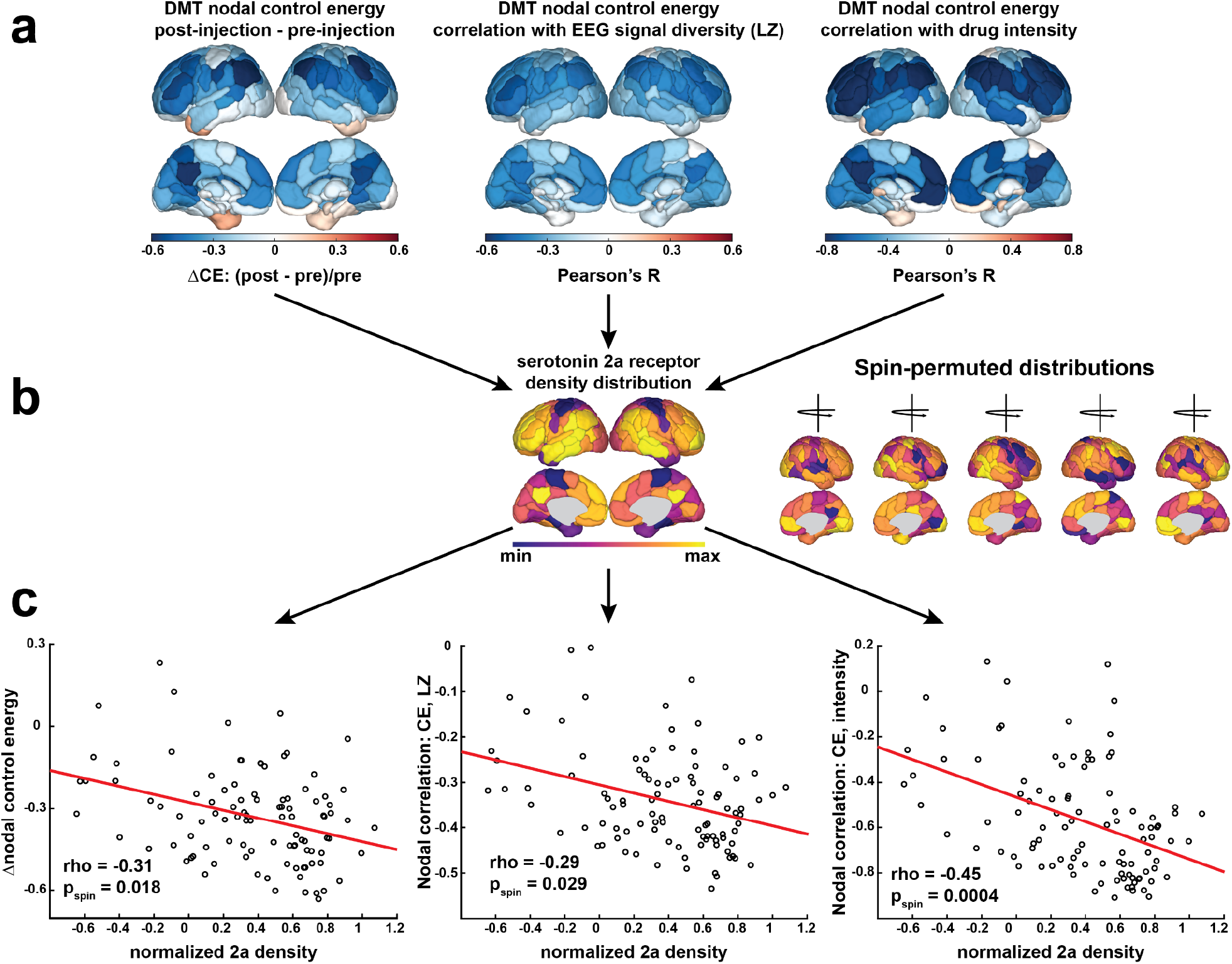
Regional control energy and its temporal correlation with signal diversity and drug intensity are associated with serotonin 2a receptor maps. **(a)** Regional control energy metrics. (left) The change in regional control energy in the 8 minutes after DMT injection, relative to the 8 minutes prior to the injection. (middle) Each region’s control energy time-series over the course of the full 28 minute DMT scans correlated with global signal diversity from EEG during the same scans. (right) Regional control energy during the DMT scans was averaged over one minute windows corresponding to the timing of subjective drug intensity ratings from separate scans. The windowed control energy time-series for each region was then correlated with the subjective drug intensity ratings. **(b)** Each of the regional metrics in (a) were then correlated with the cortical spatial map of the serotonin 2a receptor derived from PET (Beliveau et al. 2017). The strength of these correlations were compared against null correlations with 10,000 cortical spin permutations (Váša et al. 2018) of the 2a receptor map. **(c)** Scatter plots of the three cortical regions’ metrics from (a) and serotonin 2a receptor density from (b).

Through dominance analysis, we next compare the strength of the regional metrics’ associations with serotonin 2a receptor maps against their associations with other serotonin receptor subtypes, namely the serotonin (5-HT) 1a, 5-HT1b, and 5-HT4 receptors, and the serotonin transporter (5-HTT). Dominance analysis is a method of ranking the importance of multiple input variables (in our case, the five different serotonin receptor/transporters) in explaining a target variable through a series of linear regression models (Azen and Budescu 2003). A separate dominance analysis was run with each of the three metrics from Figure 3a as the target variable. We find that serotonin 2a density is the most dominant variable when explaining the variance in all three regional metrics (Figure 4).

**Figure 4:**
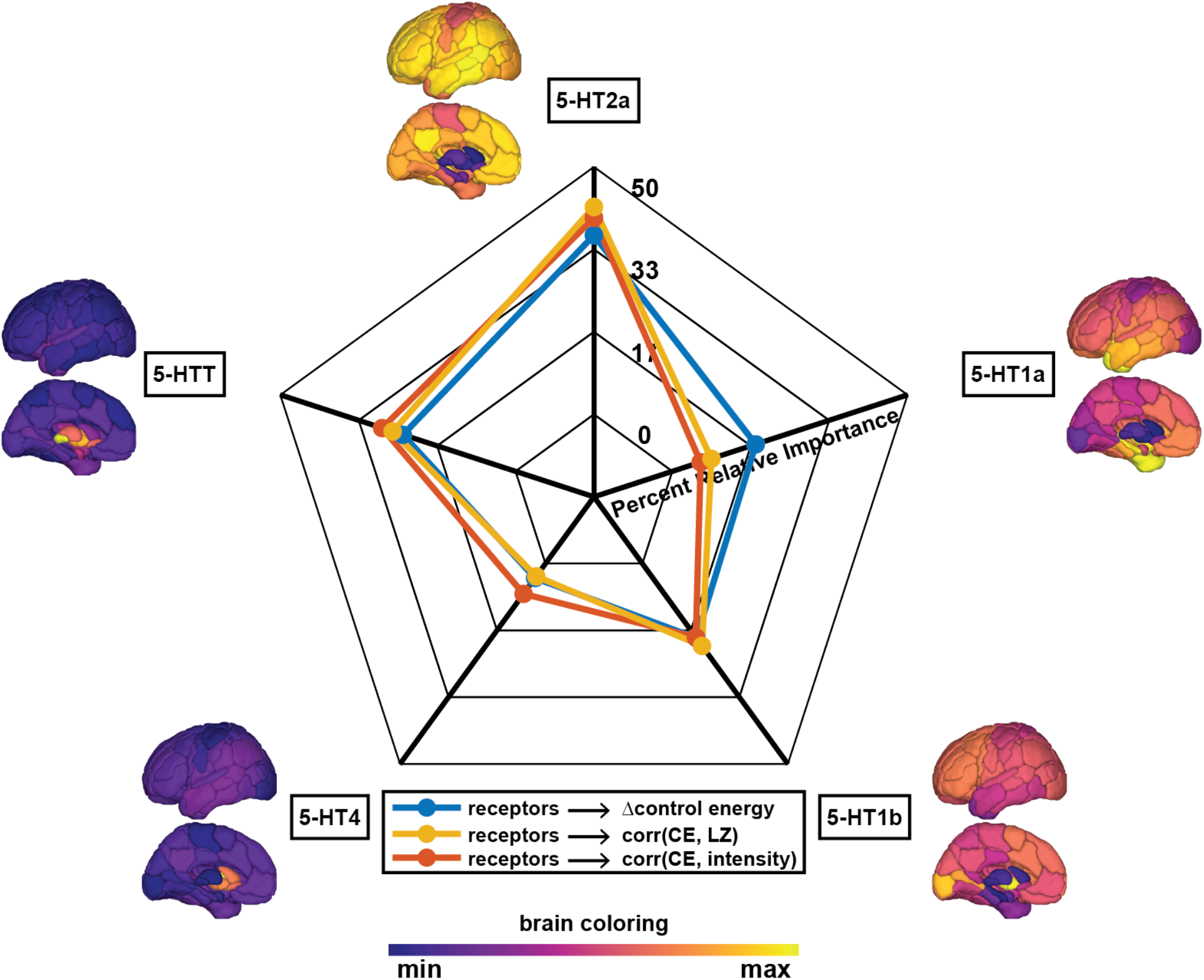
Dominance analysis reveals highest relative importance of the serotonin 2a receptor in DMT-related changes in cortical activity metrics. Three separate dominance analyses were performed using cortical values from five PET-derived serotonin receptor and transporter spatial densities (Beliveau et al. 2017) as input variables and each cortical metric from Figure 3a as the output. Dominance analysis assesses the relative importance of each input in explaining the output variable’s variance while controlling for the contributions of other predictors in multiple regressions (Azen and Budescu 2003). Displayed is the percent relative importance given to each receptor/transporter map for explaining the variance in each cortical metric, as determined by dominance analysis. 5-HT = serotonin (5-hydroxytryptamine).

### DMT’s impacts on control energy can be simulated from pharmacokinetics and the serotonin 2a receptor maps

For our simulation of DMT’s impact on control energy, we begin with all participant’s placebo fMRI scans. Prior to DMT injection, our control strategy, *B*, is uniform. Thus, pre-injection, our simulation matches the placebo control energy from Figure 2a. Following injection, we begin adding control to the system in a time and space-dependent manner according to our pharmacologically-derived time-varying control strategy (Figure 5, top). This strategy successfully estimates DMT’s impact on global control energy during the 20 minutes post-injection (Figure 5, bottom).

**Figure 5:**
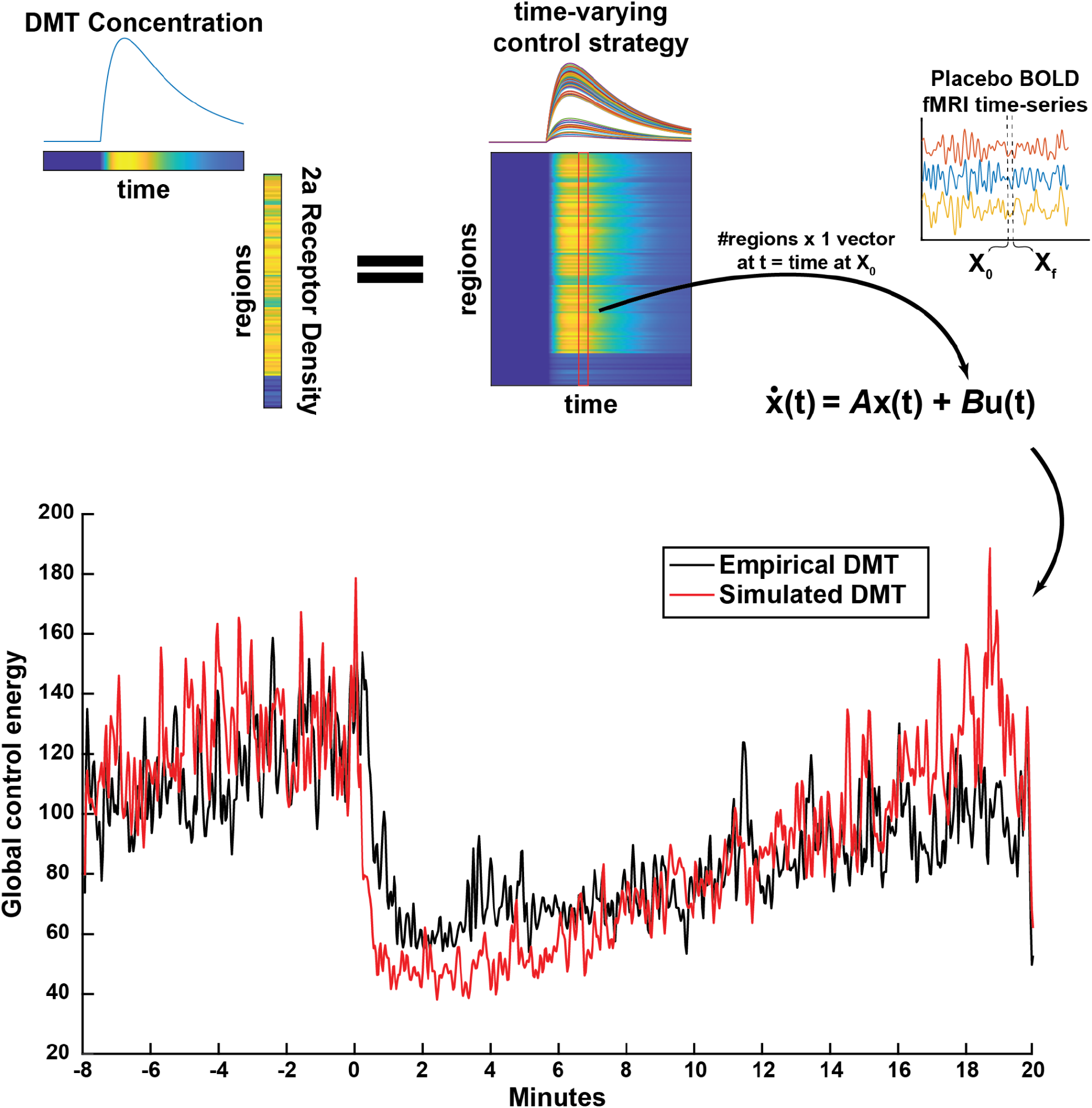
Global control energy time-series for the DMT condition is simulated using only placebo fMRI data, coupled with simulated DMT plasma concentrations and 2a receptor density information. Pharmacokinetic modeling yields an estimate of DMT concentration over the length of the 28 minute scans. Here, we specifically used predicted ‘brain-effect compartment’ concentrations from a previously validated model using plasma concentration sampling and EEG (Eckernäs et al. 2023). Multiplying DMT concentration over time by regional PET-derived serotonin 2a densities (Beliveau et al. 2017) yields an estimate for DMT’s impact on each brain region over time which can be used as a time-varying control strategy. In order to simulate the impact of DMT on the global control energy time-series, we use each participant’s placebo fMRI data and apply the time-varying control strategy via inclusion in the diagonal of matrix *B*. Prior to DMT injection, the control strategy (diagonal in *B*) is uniform, as is the case for all previously calculated energy metrics.

### Supplemental analyses

We performed several analyses to supplement our main findings. First, we calculate the time-resolved control energies for *a priori* resting-state networks (Yeo et al. 2011). We find that the most prominent reductions in control energy under DMT compared with placebo occur in the visual, frontoparietal, and default mode networks (SI Figure 1). We find that DMT’s impact on the frontoparietal and default mode network are strongest in the first ten minutes after injection. Interestingly, DMT’s impacts on the visual network are strongest in the final ten minutes after injection (SI Figure 2). Next, we reproduce our main results without the use of global signal regression during fMRI preprocessing (SI Figure 3). We find that these results are consistent with the main text, however the regional metrics are less varied and have weaker correlations with 2a receptor maps. We also show scatter plots as in Figure 3, but with subcortical regions included (SI Figure 4). The corresponding correlations are all strong and negative, however, spin testing was not performed to calculate the p-values as subcortical regions cannot be spin permuted. In SI Figure 5, we show distance metrics between true and simulated DMT control energies for two alternative models to demonstrate the advantage of incorporating both the simulated brain-effect concentration (compared to simulated plasma concentration) and the spatial 2a receptor density map (compared to uniform spatial map). Lastly, we demonstrate DMT significantly reduces global control energy under a variety of BOLD signal normalization approaches (SI Figure 6).

## Discussion

In this work, we use a time-resolved network control theory framework to characterize brain activation dynamics underlying the DMT psychedelic experience (Figure 1). We find that, on a global level, control energy is reduced under DMT compared to placebo. We then interrogate the effect of DMT in a temporally- and spatially-resolved manner. Temporally, we find that the increase in global control energy under DMT covaries with increases in EEG signal diversity and subjective drug intensity (Figure 2). Spatially, we find that the regional distribution of DMT’s effects is aligned with serotonin 2a receptor densities (Figures 3 and 4). Finally, we demonstrate a computational framework for predicting DMT’s impact on global brain dynamics using a network control model informed by pharmacokinetics and pharmacodynamics (Figure 5).

Comparing global control energies between DMT and placebo reveals an overall reduction in the input necessary for transitioning between brain-states following injection with DMT (Figure 2a). Prior work with other psychedelics (LSD and psilocybin) has demonstrated that these serotonergic 2a agonist compounds increase the diversity of brain state dynamics (Lord et al. 2019; Carhart-Harris et al. 2014; Atasoy et al. 2017; Girn et al. 2022; Luppi et al. 2021; Tagliazucchi et al. 2014; Schartner et al. 2017). Decreased control energy may be reflective of a system poised near a state of criticality, whereby lowered barriers facilitate access to a larger repertoire of state dynamics (Girn et al. 2023; Carhart-Harris and Friston 2019; Toker et al. 2022). In our own recent work using an approach similar to the one employed here, we found that LSD and psilocybin decreased control energy, and, across individuals, larger decreases under LSD were associated with more complex brain-state sequences (Singleton et al. 2022). The present study further strengthens evidence for this association between control energy and neural entropy by showing that the global control energy throughout the fMRI time-series is temporally coupled with neural signal diversity measured with simultaneous EEG (Figure 2b).

Signal diversity here refers to the Lempel-Ziv complexity of EEG signal averaged across electrodes, and its increase has been a consistent characteristic of acute psychedelic experiences (Timmermann et al. 2019; Schartner et al. 2017). We also find that control energy over time is inversely related to the intensity of the subjective drug effects (Figure 2c), linking our fMRI-based metric to participants’ experiences in real time.

We next sought to interrogate regional differences in DMT’s effects on control energy, and its association with EEG signal diversity and subjective intensity. In general, the regional metrics reflect what is observed at a global level - namely, regional control energy is decreased following injection with DMT and is inversely correlated with EEG signal diversity and subjective drug intensity (Figure 3a). Of particular interest to us was the extent to which regional heterogeneity in these metrics might be explained by biologically relevant information. The serotonin 2a receptor is the primary target in the brain responsible for initiating a cascade of changes that give rise to the characteristic subjective and neural effects of psychedelics (Kraehenmann et al. 2017; Preller, Burt, et al. 2018; Preller, Schilbach, et al. 2018). Our previous simulation studies demonstrated that the serotonin 2a receptor spatial map was particularly suited for lowering global control energies (Singleton et al. 2022). Here, we further demonstrate that regional differences in empirical control energies during psychedelic administration are related to regional serotonin 2a receptor densities (Figure 3c). We rank-correlated each of the three regional metrics’ values with that of the serotonin 2a cortical distribution. In each case, we find that the regional spatial pattern of control energy differences and its association with signal diversity and drug intensity are inversely related to the density of the serotonin 2a receptor, above and beyond the effect of spatial autocorrelation (Figure 3c).

One might ask, however, whether the serotonin 2a receptor is associated with the regional metrics at a level above and beyond other serotonin system receptors. To answer this question, we performed a dominance analysis (Azen and Budescu 2003) for each regional metric using five input variables: the serotonin 2a (5-HT2a), serotonin 1a (5-HT1a), serotonin 1b (5-HT1b), serotonin 4 (5-HT4) receptors, and the serotonin transporter (5-HTT). The serotonin 2a receptor was found to have the highest relative importance in explaining the variance of regional decreases in control energy after DMT, as well as the control energy’s correlation with signal diversity and subjective intensity (Figure 4). Together, these results suggest that regions having higher densities of the serotonin 2a receptor are the most impacted by DMT, and have stronger couplings with neural and subjective effects.

Having established an association between DMT’s impact on control energy and the serotonin 2a receptor distribution, we finally ask the question: “can DMT’s impact on control energy be simulated from non-drug data (i.e., in this case, the placebo dataset) using a pharmacologically-informed control framework”? Our control energy calculations up to this point were agnostic to regional heterogeneity, i.e. they deployed a uniform control strategy (encoded by the control matrix *B* being the identity). However, adjustments to the control strategy have been successfully deployed for *in silico* hypothesis testing (Cornblath et al. 2020; He et al. 2022; Zhou et al. 2021; Parkes et al. 2022) and simulations of external and internal forms of stimulation (Singleton et al. 2022; Stiso et al. 2019; Luppi et al. 2023). DMT injection exhibits rapid onset of subjective and neural effects in a concentration-dependent manner (Strassman et al. 1994; Strassman 1995). Previous work has demonstrated successful pharmacokinetic modeling of DMT plasma concentration and its neural effects at dosing regimens similar to those used in the present study (Eckernäs et al. 2022; 2023). Here, we used an independently validated model of DMT’s pharmacokinetic impact on EEG alpha rhythms to simulate population-level ‘brain-effect’ concentrations of DMT over the course of the 28 minute scans (Eckernäs et al. 2023). We next combine this temporal estimation of DMT’s effects with spatial information by multiplying the simulated DMT concentration at each time-point with the regional serotonin 2a receptor maps (Beliveau et al. 2017). This yields a temporal and spatial map of DMT’s hypothesized impact on brain dynamics, which we operationalize as a time-varying control strategy within our network control theory approach (Figure 5). Our ability to estimate DMT’s impacts on control energy through pharmacologically-informed adjustments to model parameters serves as an important proof-of-concept for using network control theory in computational psychiatry applications (Moujaes et al. 2022; Vohryzek et al. 2023). Importantly, we validated the superior utility of our two specific pieces of information in the control strategy - the ‘brain-effect’ versus the plasma concentration and the 2a receptor’s spatial information versus a uniform control approach (SI Figure 4).

The present work validates a meaningful fMRI correlate of the increased signal complexity or entropy that has been reliably demonstrated with EEG or MEG recordings of the psychedelic state. The latter modalities have limited spatial resolution (EEG) or depth (MEG), precluding inferences about specific regional effects or changes in deep structures. Thus, observing fMRI correlates of other modalities’ recordings deepens our understanding of the so-called ‘entropic brain’ effect (Carhart-Harris et al. 2014; Carhart-Harris 2018). The strength of fMRI is its high spatial resolution and whole-brain coverage; however, relative to EEG, its cost, immobility and other practical challenges limit its widespread application. Recent work has shown that EEG-recorded signal complexity or entropy can be effectively tracked in real time (Mediano et al. 2023), inspiring ideas regarding how such information could be used to adapt treatment parameters such as dosage, in a way to suit individual responses. More work across all modalities is required to further deepen our understanding of mechanisms of psychedelic action, e.g., to test whether the observed acute brain effects begin with a 5-HT2a receptor agonism initiated spike-to-wave decoupling (Celada et al. 2008) translating downstream into anatomical and functional neuroplastic changes such as those found in preclinical (Ly et al. 2018; Shao et al. 2021; Hesselgrave et al. 2021; Cameron et al. 2023; Vargas et al. 2023) and clinical neuroscience research (Doss, Považan, et al. 2021; Daws et al. 2022).

Although the present analysis expands on prior work in this field, several important limitations must be considered. First, our small sample size limits the generalizability of our findings; however, this is the third independent dataset which has found decreased control energy during psychedelic administration, which lends more confidence in our findings (Singleton et al. 2022). Our time-resolved control energy analysis provides the ability to associate control energy changes in real time with other imaging metrics and subjective effects. However, like many time-resolved metrics, this increases noise sensitivity. For our correlational analyses, we consider associations between group-level metrics in order to more accurately assess changes over time and space. In addition, we assess our simulation’s success based on the group-average outputs from our model. Extending the work towards individual level modeling will be a fruitful avenue for future research, with implications for personalized medicine (Moujaes et al. 2022; Vohryzek et al. 2023). Elucidating the potential impacts of vasoconstriction (Gamoh, Hisa, and Yamamoto 2013) and arousal (Liu and Falahpour 2020) on our findings also requires sophisticated experimental designs (e.g., using an active stimulant as control) and should be investigated in future studies.

In summary, we have demonstrated that time-resolved network control analysis captures meaningful changes in brain activation dynamics during the onset, peak, and offset of an infused DMT experience. We found significant reductions in the control energy required for the brain to traverse its activity landscape under DMT, and an association between decreases in control energy and increases in EEG-based neural signal complexity and subjective drug effects in a manner that is regionally related to serotonin 2a receptor density. Finally, we demonstrated that through a pharmacologically-informed network control framework, that we are able to simulate DMT’s impacts on control energy over time - an important step towards understanding mechanisms of these neuromodulators.

## Methods

### Participants and Experimental Procedures

The original single blind, placebo controlled, counterbalanced study was approved by the National Research Ethics (NRES) Committee London – Brent and the Health Research Authority and was conducted under the guidelines of the revised Declaration of Helsinski (2000), the International Committee on Harmonisation Good Clinical Practices guidelines, and the National Health Service Research Governance Framework. Imperial College London sponsored the research, which was conducted under a Home Office license for research with Schedule 1 drugs.

The original study is presented in (Timmermann et al. 2023) however we summarize the relevant design material here. Volunteers participated in two testing days, separated by 2 weeks. On each testing day, participants arrived and were tested for drugs of abuse and were involved in 2 separate scanning sessions. In this initial session (task-free) they received intravenous (IV) administration of either placebo (saline) or 20 mg DMT (in fumarate form) in a counterbalanced order (half of the participants received placebo and the other half received DMT). This first session always consisted of continuous resting-state scans which lasted 28 minutes with DMT/placebo administered at the end of 8^th^ minute. Participants laid in the scanner with their eyes closed (an eye mask was used to prevent eyes from opening), while EEG activity was recorded. A second session then followed with the same procedure as the initial session, except on this occasion participants were asked to rate the subjective intensity of drug effects every minute in real time. The design was single blind (only researchers were aware of the order of administration).

This article only analyzes the resting-state scans in which no intensity ratings were asked, but uses the intensity ratings for correlational analyses. In total, 20 participants completed all study visits (7 female, mean age = 33.5 years, SD = 7.9).

### fMRI and EEG acquisition

The original study is presented in (Timmermann et al. 2023), however we summarize the relevant acquisition information here. Images were acquired in a 3T MRI (Siemens Magnetom Verio syngo MR B17) using a 12-channel head coil for compatibility with EEG acquisition.

Functional imaging was performed using a T2*-weighted BOLD sensitive gradient echo planar imaging sequence with the following parameters: repetition time (TR) = 2000ms, echo time (TE) = 30ms, acquisition time (TA) = 28.06 mins, flip angle (FA) = 80°, voxel size = 3.0 x 3.0 x 3.0mm3, 35 slices, interslice distance = 0mm. Whole-brain T1-weighted structural images were also acquired.

EEG was recorded inside the MRI during image acquisition at 31 scalp sites following the 10-20 convention with an MR compatible BrainAmp MR amplifier (BrainProducts, Munich, Germany) and an MR-compatible cap (BrainCap MR; BrainProducts GmbH, Munich, Germany). Two additional ECG channels were used to improve heart rate acquisition for artifact minimization during EEG preprocessing. EEG was sampled at 5 kHz and with a hardware 250 Hz low-pass filter. EEG-MR clock synchronization was ensured using the Brain Products SyncBox hardware.

### fMRI preprocessing

The same preprocessing pipeline as used in previous work with LSD and psilocybin (Singleton et al. 2022) and reported in Timmermann and colleagues (2023) was used here. Six out of 20 participants were discarded from group analyses due to excessive head movement during the 28 minute DMT scans (>15% of scrubbed volumes with a scrubbing threshold of frame-wise displacement (FD) of 0.4 (Power et al. 2014)), leaving 14 for analysis. Preprocessing steps consisted of 1) de-spiking (3dDespike, AFNI (Cox 1996)), 2) slice time correction (3dTshift, AFNI), 3) motion correction (3dvolreg, AFNI) by registering each volume to the most similar volume, in the least squares sense, to all others (in-house code), 4) brain extraction (BET, FSL (Smith et al. 2004)), 5) rigid body registration to anatomical scans, 6) non-linear registration to 2mm MNI brain (Symmetric Normalization (SyN), ANTS (Avants, Tustison, and Song 2009)), 7) scrubbing - using an FD threshold of 0.4 - with scrubbed volumes being replaced with the mean of the surrounding volumes. Additional preprocessing steps included: 8) spatial smoothing (FWHM) of 6mm (3dBlurInMask, AFNI), 9) band-pass filtering between 0.01 to 0.08 Hz (3dFourier, AFNI), 10) linear and quadratic de-trending (3dDetrend, AFNI), 11) regressing out 9 nuisance regressors (all nuisance regressors were bandpassed filtered with the same filter as in step 9), out of these, 6 were motion-related (3 translations, 3 rotations) and 3 were anatomically-related. Lastly, global signal regression was performed and time-series were parcellated into 100 cortical (Schaefer et al. 2018) and 16 subcortical (Tian et al. 2020) regions of interest.

### Structural connectivity network construction

The structural connectome used for network control theory analysis was identical to the one used in prior work (Luppi et al. 2021; Singleton et al. 2022). Namely, we relied on diffusion data from the Human Connectome Project (HCP, http://www.humanconnectome.org/), specifically from 1021 subjects in the 1200-subject release (Van Essen et al. 2013). A population-average structural connectome was constructed and made publicly available by Yeh and colleagues (2018) in the following way. Multishell diffusion MRI was acquired using b-values of 1000, 2000, 3000 s/mm^2^, each with 90 directions and 1.25 mm iso-voxel resolution. Following previous work (Luppi et al. 2021; Yeh et al. 2013; Singleton et al. 2022), we used the QSDR algorithm implemented in DSI Studio (http://dsi-studio.labsolver.org) to coregister the diffusion data to MNI space, using previously adopted parameters (Yeh et al. 2013). Deterministic tractography with DSI Studio’s modified FACT algorithm then generated 1,000,000 streamlines, using the same parameters as in prior work, specifically, angular cutoff of 55?, step size of 1.0 mm, minimum length of 10 mm, maximum length of 400mm, spin density function smoothing of 0.0, and a QA threshold determined by DWI signal in the CSF. Each of the streamlines generated was screened for its termination location using an automatically generated white matter mask, to eliminate streamlines with premature termination in the white matter. Entries in the structural connectome *A*_*ij*_ were constructed by counting the number of streamlines connecting every pair of regions *i* and *j* in the augmented Schaefer-116 atlas (Schaefer et al. 2018; Tian et al. 2020). Lastly, streamline count was normalized by the number of voxels contained in each pair of regions.

### 5-HT receptor mapping

Details for obtaining the serotonin receptor density distribution have been previously described (Beliveau et al. 2017) however we provide a brief summary here. PET data for 210 participants (not under the influence of psychedelics) were acquired on a Siemens HRRT scanner operating in 3D acquisition mode with an approximate in-plane resolution of 2mm (1.4 mm in the center of the field of view and 2.4 mm in cortex) (Olesen et al. 2009). Scan time and frame length were designed according to the radiotracer characteristics. For details on MRI acquisition parameters, which were used to coregister the data to a common atlas, see Knudsen et al (2016). The voxelwise average density (Bmax) maps for each receptor were parcellated into 116 regions of interest for the augmented Schaefer-116 atlas (Schaefer et al. 2018; Tian et al. 2020).

### EEG signal diversity: Lempel-Ziv complexity

Following our previous study involving DMT (Timmermann et al. 2019), as well as those performed with LSD, psilocybin and ketamine (Schartner et al. 2017), we performed signal diversity analysis using the Lempel-Ziv 1976 algorithm (LZ76), as reported in Timmermann et al. (2023). The EEG signal at each single electrode was binarized using its mean for each 2-second epoch, and then the LZ76 algorithm was used to generate a dictionary of unique subsequences whose size quantifies the temporal diversity for the signal (denoted here as LZs). The average LZs across channels was used for correlational analyses with control energy.

### Dominance Analysis

Dominance analysis was used in order to determine the relative importance of each receptor/transporter map on predicting nodal control energy metrics. Dominance analysis aims to establish the relative significance (or “dominance”) of each independent variable in relation to the overall fit (adjusted R^2^) of the multiple linear regression model (https://github.com/dominance-analysis/dominance-analysis) (Azen and Budescu 2003). This process involves fitting the same regression model to every possible combination of predictors (creating 2^p^ – 1 submodels for a model with p predictors). Total dominance is characterized as the mean increase in R^2^ when incorporating a single predictor of interest to a submodel, considering all 2^p^ – 1 submodels. The aggregate dominance of all input variables equals the total adjusted R^2^ of the comprehensive model, rendering the relative importance percentage an easily understandable technique that apportions the overall effect size among predictors. As a result, in contrast to alternative methods for evaluating predictor significance, such as those based on regression coefficients or univariate correlations, dominance analysis takes into account predictor-predictor interactions and offers interpretability.

### Minimum control energy

Network control theory allows us to probe the constraints of white-matter connectivity on dynamic brain activity, and to calculate the minimum energy required for the brain to transition from one activation pattern to another (Singleton et al. 2022; Cornblath et al. 2020; Karrer et al. 2020). While this procedure has been detailed elsewhere (Karrer et al. 2020), we summarize it briefly here. We obtained a representative NxN structural connectome *A* obtained as described above using deterministic tractography from HCP subjects (*see Methods and Materials; Structural Connectivity Network Construction*), where N is the number of regions in our atlas. We then employ a linear time-invariant model:

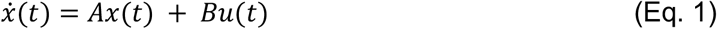

where x is a vector of length N containing the regional activity at time *t. B* is an NxN matrix that contains the control input weights, and is otherwise known as the control strategy. In our analyses, *B* is constructed by placing an input vector, *v*, along the diagonal of the matrix *B*. In uniform control scenarios, *v* is a vector of length N containing all ones. In the case of our DMT simulation, a time-varying control strategy (*B*) was used (Figure 5, top), where the input vector, *v* was a function over time of simulated DMT concentration *DMT*(*t*), the regional serotonin 2a receptor density vector ρ, and a scaling parameter α:

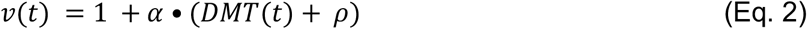

which effectively adds additional control to the system as a function of increasing DMT, in a manner that is regionally skewed by 2a density. In order to estimate the scaling parameter α, we performed a grid search over values [1, 10, 20, 30, 40, 50, 60, 70] and chose the value which minimized the Euclidean distance between the simulated output and the empirical DMT control energy on a group-level (SI Figure 4; alpha = 30). DMT concentration over time was simulated from previously published population level pharmacokinetic parameter estimates to obtain the typical predicted concentrations after a bolus dose of 20 mg DMT fumarate (using the R package *mrgsolve*) (Eckernäs et al. 2023). Specifically, theoretical effect compartment concentrations (based on EEG Alpha rhythms) were operationalized as *DMT(t)*, as this can be thought of as representing DMT concentration in the brain. The regional serotonin 2a receptor density ρ was derived from previously published PET data (*See Methods: 5-HT receptor mapping*) (Beliveau et al. 2017).

To compute the minimum control energy required to drive the system (network) from an initial activity pattern (x_0_) to a final activity pattern (x_f_) over some finite amount of time (*T*), we minimize the inputs (*u(t)*) subject to Equation 1:

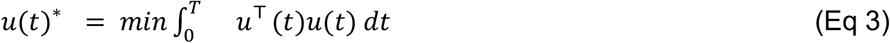

where *T* is the time horizon that specifies the time over which input to the system is allowed. Here, a common choice of *T* = 1 was used (Braun et al. 2021; Betzel et al. 2016; Parkes et al. 2022; Karrer et al. 2020). The minimum control energy for a single brain region *i* is then:

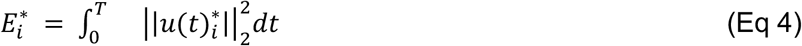

And, finally, the global minimum control energy for a transition is the sum of Eq 4 over each region:

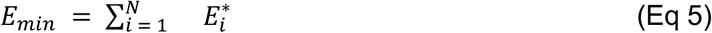

This quantity was calculated for each pair of initial x_0_ and final x_f_ brain states (i.e. adjacent BOLD volumes in each individual’s fMRI scans) to obtain a time-series of control energy over each individuals’ 28 minute scanning sessions.

## Data and code availability

Data used in this analysis were published alongside the original study (Timmermann et al. 2023). Code to reproduce this analysis is available at: https://github.com/singlesp/DMT_NCT.

## Acknowledgements

This work was funded in part by NIH grants: R01NS102646 (AK) and RF1MH123232 (AK). AIL is supported by the Molson Neuro-Engineering Fellowship and FRQNT Strategic Clusters Program (2020-RS4-265502 - Centre UNIQUE - Union Neuroscience & Artificial Intelligence - Quebec) via the UNIQUE Neuro-AI Excellence Scholarship. The original DMT EEG-fMRI study data collection was made possible by donations from Patrick Vernon, mediated by the Beckley Foundation, as well as supplementary support from Anton Bilton and other founders of the Centre for Psychedelic Research, Imperial College London. CT is supported by funders for the Centre for Psychedelic Research at Imperial College London. RCH is supported by the Ralph Metzner Chair in Neurology and Psychiatry at UCSF.

