## Supplemental_Information for "Time-resolved network control analysis links reduced control energy under DMT with the serotonin 2a receptor, signal diversity, and subjective experience"

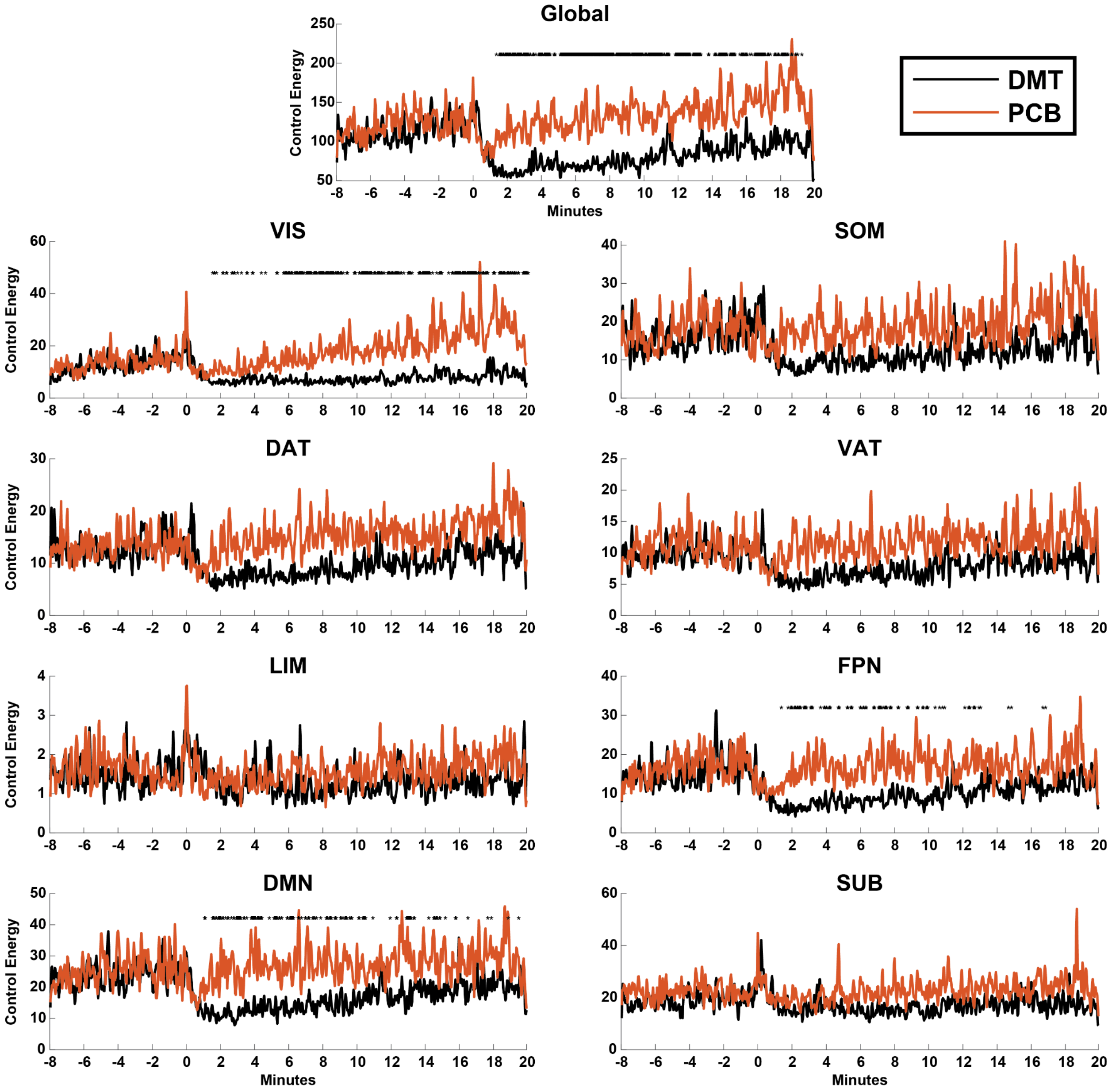


**SI Figure 1: Subnetwork level comparisons in post-injection control energy.** Here, we replicate the main text results from Figure 2a. At the top, we reprint the global results from Figure 2a for the reader's convenience. Below, we repeat the analysis for each of the well-characterized intrinsic connectivity subnetworks of Yeo and colleagues (2011), plus an additional subcortical one. The most prominent reductions in control energy occur in the visual, frontoparietal, and default mode (sub)networks. Each subnetwork's control energy was obtained by summing control energy over all nodes assigned to that network. Control energies for each transition following injection were compared via a two-sided, paired t-test and p-values were corrected for multiple comparisons using the Benjamini-Hochberg method (* = corrected p <

0.05). VIS = visual; SOM = somatomotor; DAT = dorsal attention; VAT = ventral attention; LIM = limbic; FPN = frontoparietal; DMN = default mode; SUB; subcortex.

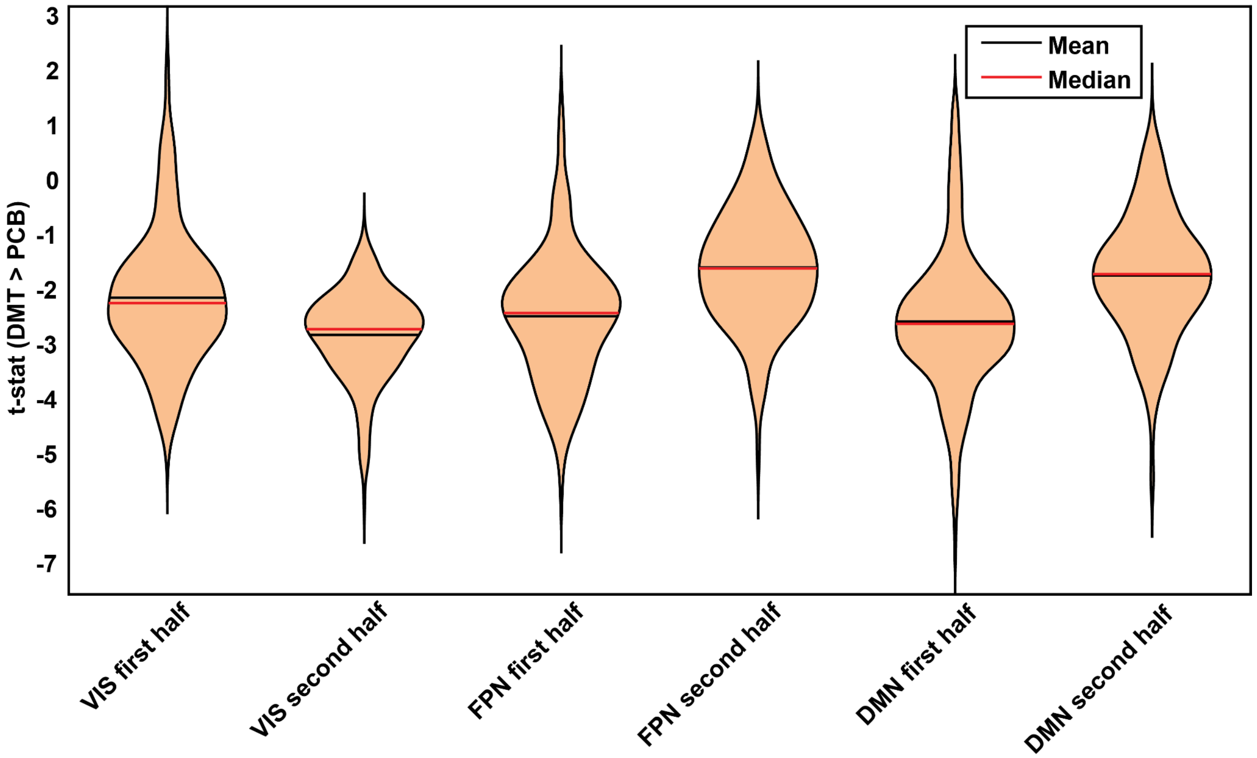


**SI Figure 2: Subnetwork control energy differences in the first and second half of post-injection scanning.** For each subnetwork that was significantly impacted by DMT (SI Figure 1), we plot the average t-statistic for transitions occurring in the first ten minutes after injection (first half) and the last ten minutes after injection (second half). We note that the frontoparietal (FPN) and default mode networks (DMN) are most greatly impacted in the first ten minutes after injection, whereas the opposite is true for the visual network (VIS).

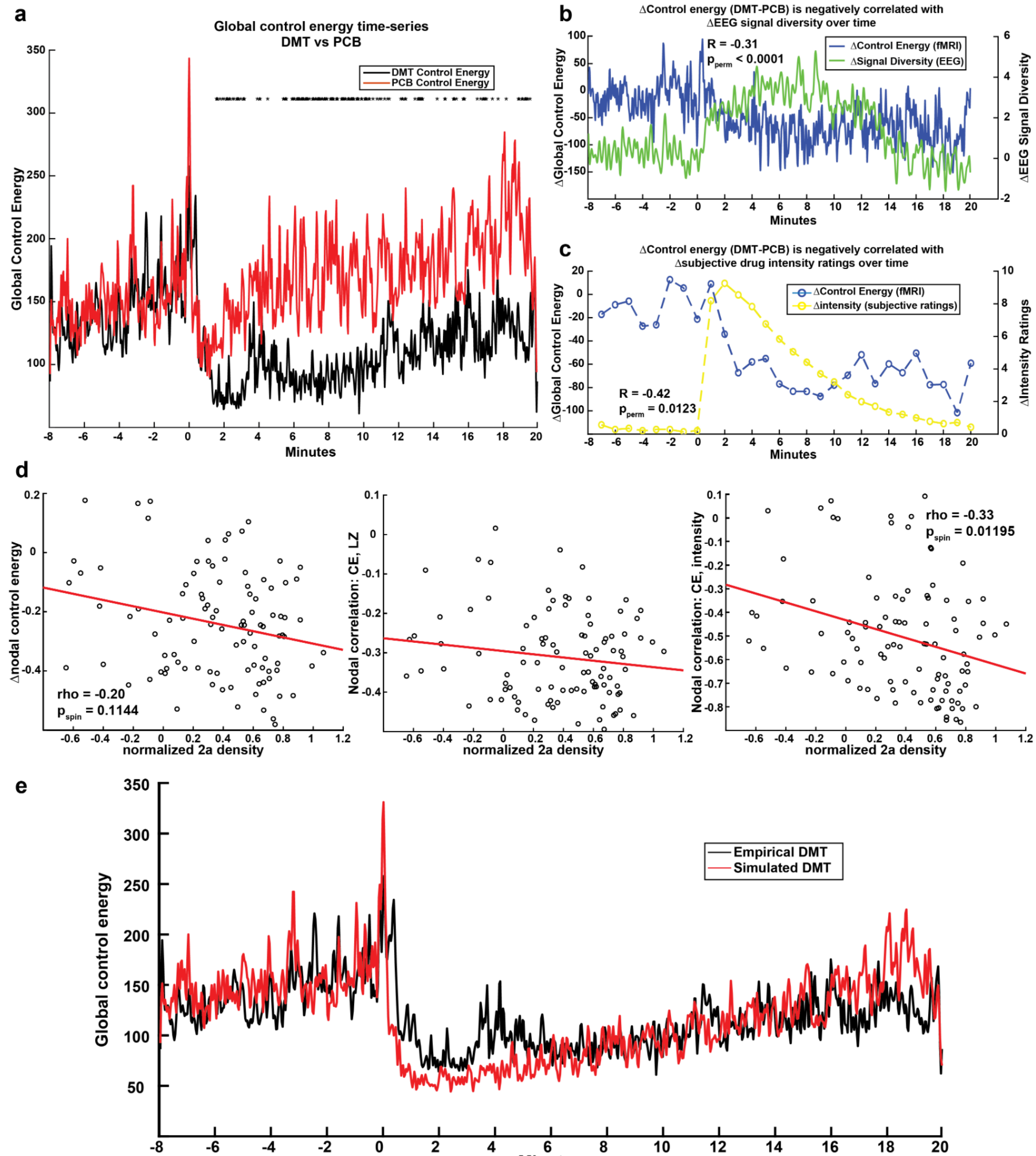


**SI Figure 3: Replication of main results without the use of global signal regression.** (a-c) Main results from Figure 2. (d) Main results from Figure 3c. (e) Main results from Figure 5.

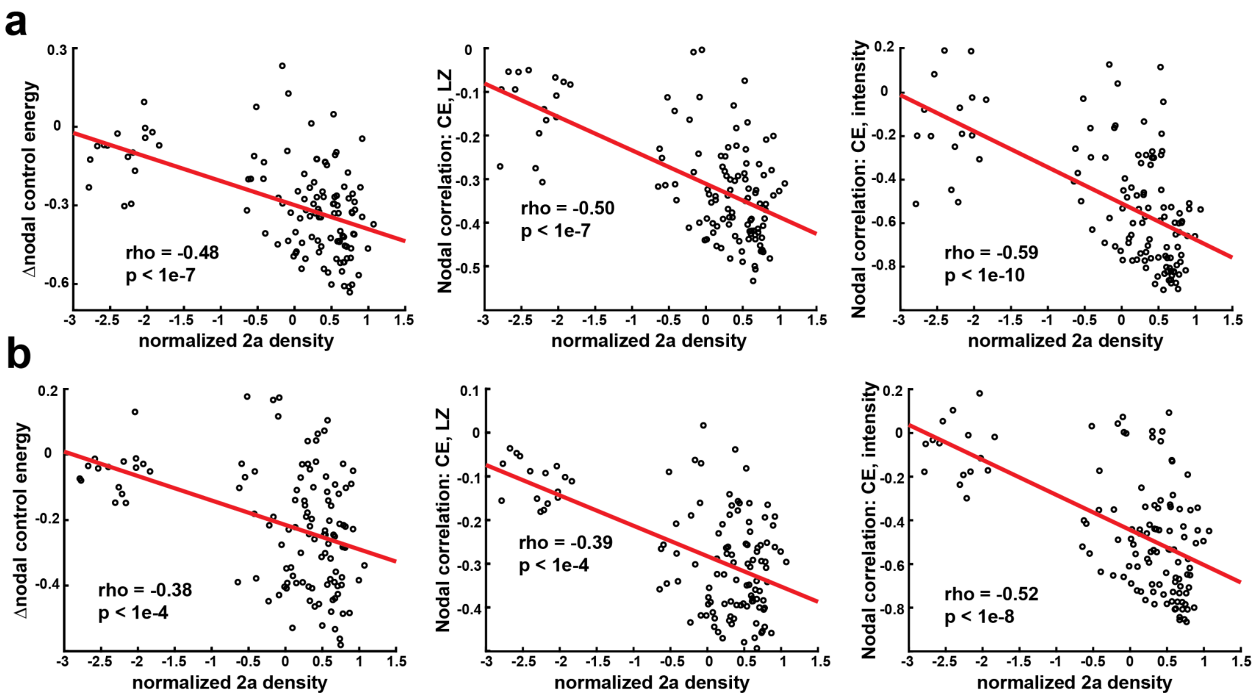


**SI FIgure 4: Nodal metric correlations including the subcortex.** (a) Scatter plots from Main Figure 3c (no global signal regression), repeated with the subcortical regions included. (b) Scatter plots from SI Figure 2d (with global signal regression), repeated with the subcortical regions included. Spearman rank correlations, uncorrected p-values.

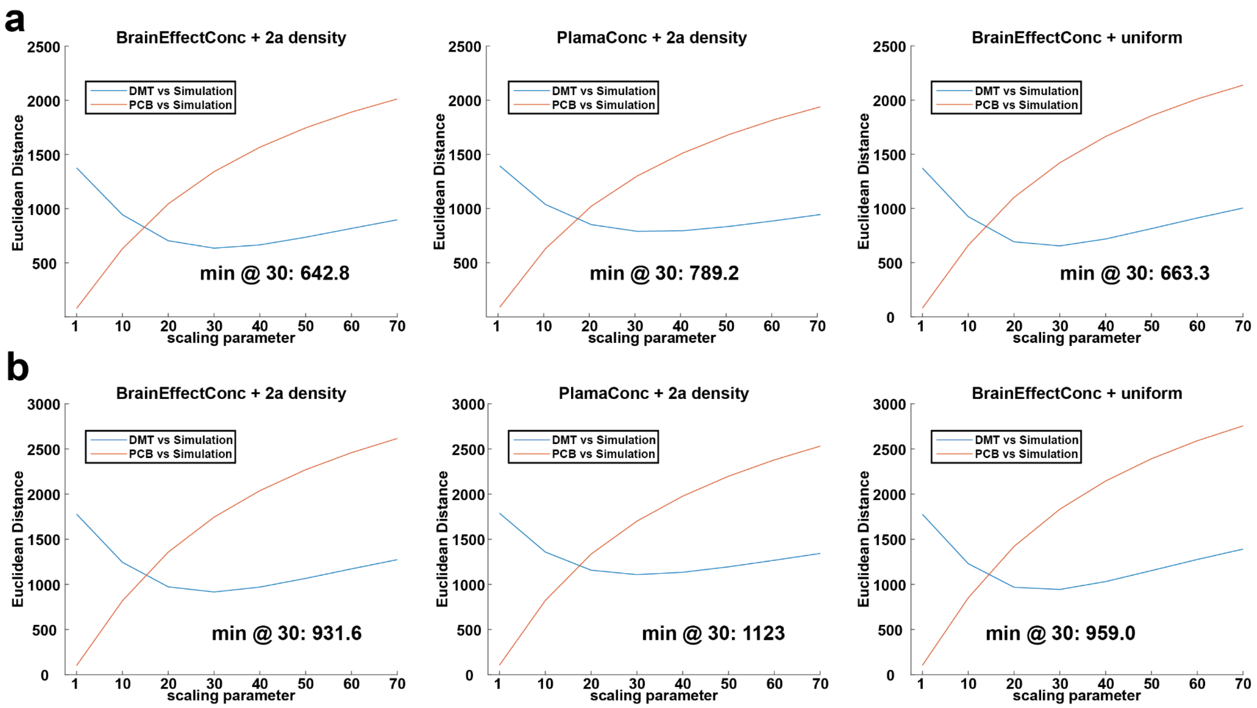


**SI Figure 5: Optimization of scaling parameter for model simulations.** (a) Main text simulations (with global signal regression). (b) Simulations without the use of global signal regression. (left) Optimization of scaling parameter for the main model presented in the main text, which uses simulated brain effect compartment concentrations for DMT's impact over time, and the serotonin 2a density for spatial differences. (middle) The first comparison model which uses simulated plasma concentration rather than brain effect concentration. (right) The second comparison model, which removes the effect of the serotonin 2a spatial map in the original model by adding uniform control in its place. In this second model, the same amount of overall control is given to the system as in the original simulation. We note the Euclidean distance quantifying model error is minimal in the first model's simulations for both sets of data.

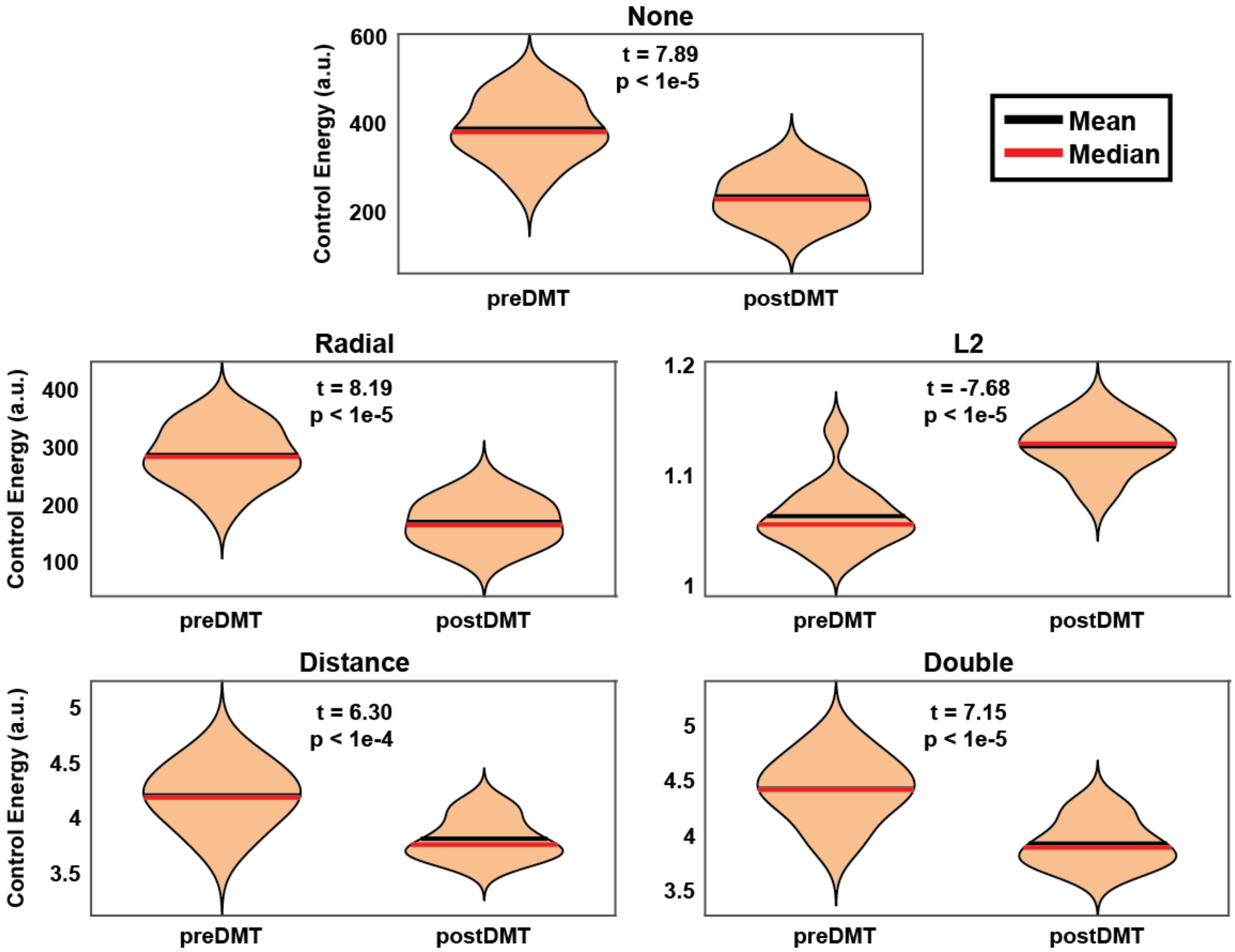


**SI Figure 6: Comparison of average global control energy over the 8 minutes prior to injection to the 8 minutes following injection, using different methods of brain activity normalization.** All methods of activity normalization result in a significant decrease in control energy after DMT injection, except for L2 normalization, which results in a significant increase in control energy. This suggests that when accounting for the magnitude of activity, states are more difficult to reach through a network control process. This is likely related to the concurrent psychedelic effect of decreased signal magnitude and increased temporal entropy. When magnitude is normalized and its effects removed from the energy calculations, the decreased autocorrelation between adjacent BOLD activity dominates the energy calculations and results in increased TE. When both magnitude and inter-state distance are accounted for (double normalization), DMT still decreases control energy. This suggests that when both factors are accounted for, states are still closer together through a network diffusion process after DMT compared to before DMT. None = no additional state normalization applied beyond fMRI preprocessing (main text version). Radial = final states are normalized so that they are unit distance from initial states. L2 = all states are normalized by their L2 magnitudes. Distance = each pair of initial and final states are normalized by the magnitude of their inter-state distance. Double = first L2 normalization is applied to all states, then distance normalization is applied to all pairs of initial and final states. a.u. = arbitrary units.
